# Local adaptation of *Gladiolus carneus* to soil nutrient extremes in the Cape Floristic Region

**DOI:** 10.1101/2025.08.01.668149

**Authors:** Katharine L. Khoury, Ethan L. Newman

## Abstract

**Premise:** Within the hyper-diverse Cape Floristic Region (CFR), divergent edaphic niches are hypothesised as an important driver of lineage diversification. However, the evidence for local adaptation on contrasting soils in this region remains limited. Using a reciprocal translocation and common garden approach, we use the polymorphic geophyte *Gladiolus carneus* to test for local adaptation to soil properties along an elevational gradient.

**Methods:** We first quantified the edaphic niche of all ecotypes of *G. carneus* across its entire range and for three experimental populations used for reciprocal translocations. Next, under common garden conditions, we tested whether the three focal populations had a fitness advantage on their native soils relative to other populations. We then assessed whether those same populations showed evidence of local adaptation.

**Key results:** We found that *G. carneus* ecotypes occupied distinct edaphic niches and that our three experimental populations occurred in nutrient-poor, intermediate-nutrient and nutrient-rich soil conditions. The common garden showed that seedlings native to the nutrient-rich and nutrient-poor sites, experienced a fitness advantage over non-native seedlings. Additionally, we found evidence of local adaptation at all translocation sites. Together, the common garden and reciprocal translocation suggests that edaphic conditions are contributing to local adaptation at the nutrient-rich and nutrient-poor sites.

**Conclusions:** We provide the first evidence of local adaptation to edaphic niches in the hyper-diverse Cape Floristic Region. These results suggest that edaphic conditions, along with other abiotic factors, may be important in driving incipient speciation in the CFR.

## INTRODUCTION

Local adaptation is commonly found in plants (Hereford, 2009), and is thought to play a key role mediating incipient speciation and lineage formation (Nosil et al., 2005; Butlin and Faria, 2024). Local adaptation evolves when heterogenous environments cause divergent selection on functional traits, giving a population a fitness advantage in their local habitat relative to foreign populations [i.e., Kawecki and Ebert’s (2004) “local versus foreign” criterion for local adaptation]. Some of the earliest studies documenting local adaptation used common garden experiments to showcase genetically based morphological differences, from different populations of the same species (*see* Turesson, 1922). Since then, local adaptation has been documented extensively between taxa occupying distinct climatic (Macel et al., 2007; Ellis and Ågren, 2024), pollinator (Newman et al., 2015; Gross et al., 2023; Johnson, 2025), coastal (Lowry et al., 2008; Popovic and Lowry, 2020), and edaphic niches (Ellis and Weis, 2006; Macel et al., 2007; Thiergart et al., 2020; Dittmar and Schemske, 2023).

As the edaphic environment can influence a plants’ access to nutrients, microbiota, and water, local adaptation to contrasting edaphic conditions has been tested extensively (*see* Rajakaruna, 2017). In particular, local adaptation to extreme edaphic conditions have received much attention, and has been tested in geothermal soils (Lekberg et al., 2012), serpentine soils (Sambatti and Rice, 2006; Yost et al., 2012; Dittmar and Schemske, 2023), and soils containing heavy metals (Jimenez-Ambriz et al., 2007). Reciprocal translocations between populations characterized by extreme and non-extreme soils, often show that both native and introduced genotypes have a higher fitness on the non-extreme, often resource-rich soils in comparison to the extreme soils (Dittmar and Schemske, 2023). Nevertheless, the increased fitness of both the native and introduced genotypes on the non-extreme soils may still fulfil the ‘local vs foreign’ criterion for local adaptation (Kawecki and Ebert, 2004), when the fitness of the native individuals are higher, than those which are introduced. This is the pattern we would expect when comparing genotypes that are suspected to be adapted to nutrient-poor and nutrient-rich soils. For example, Dittmar and Schemske (2023) found that *Leptosiphon parviflorus* on sandstone soils exhibited a home-site advantage in two of the four years tested, whereas nutrient-extreme serpentine genotypes showed a home-site advantage in all four years, albeit with lower overall fitness. Although there is good evidence for local adaptation between extreme edaphic environments (Sambatti and Rice, 2006; Jimenez-Ambriz et al., 2007; Dittmar and Schemske, 2023), the evidence on more moderate edaphic gradients is mixed (Yost et al., 2012; Jimenez-Ramirez et al., 2023; Ellis and Ågren, 2024).

Neither Jimenez-Ramirez et al. (2023) or Ellis and Ågren (2024), who both tested for local adaptation between moderate edaphic environments, found any evidence of local adaptation to soil types. However, Yost et al. (2012), found local adaptation between cryptic *Lasthenia* species on a gradient of serpentine soils. These studies demonstrate heterogenous edaphic conditions do not always result in local adaptation, and a combination of multi-year reciprocal translocations and common garden experiments are needed to comprehensively test for local adaptation to soil types (Kawecki and Ebert, 2004).

Edaphic factors are hypothesised to be a major driver of diversification in the Cape Floristic Region (CFR), however the extent of its contribution has long been questioned (Linder, 2003; Ellis et al., 2014). This is mainly due to the dearth of experimental evidence for divergent soil types driving diversification in the CFR (*but see* Verboom et al., 2004; Verboom et al., 2012), at least in comparison to phylogenetic evidence, of which the evidence is conflicting in its support (*see* van der Niet and Johnson, 2009; Schnitzler et al., 2011; Forest et al., 2014; Verboom et al., 2017). To our knowledge, no study exists that isolates local adaptation to soil conditions in the CFR. Although most soils in the CFR are highly leached and infertile, complex geological and geomorphic processes have created heterogenous soils ranging from young, relatively fertile soils, to old, extremely nutrient-poor soils (Cramer et al., 2014).

Specifically, the erosion-resistant sandstone of the Cape Fold Mountains has the most nutrient-poor soils recorded on earth (Cramer et al., 2014), with particularly low phosphorus concentrations (Linder, 2003; Cramer et al., 2014). However, the soils in the coastal planes and intermontane valleys are generally derived from shale and are moderately fertile (Cramer et al., 2014). This has resulted in a diverse array of soil niches and selective regimes that may drive divergence between closely related taxa in the CFR.

*Gladiolus carneus* (Iridaceae) is a polymorphic geophyte endemic to the CFR (Goldblatt and Manning, 2020), consisting of seven morphologically distinct ecotypes (Khoury et al., in press). These ecotypes occur along an elevational gradient and likely occupy distinct soil niches, allowing for comprehensive tests of local adaptation to divergent soil properties. Due to the heterogeneity of soil types in the CFR, and the wide geographic range of the species, we would expect *G. carneus* ecotypes to occupy a range of nutrient-rich to nutrient-poor edaphic niches. We further predict all tested ecotypes will experience increased fitness on nutrient-rich soils, and that ecotypes occupying any extreme edaphic environments (both nutrient-rich and nutrient-poor) will be locally adapted. However, ecotypes occupying less extreme edaphic niches are not expected to be locally adapted to soil type exclusively. Overall, we use environmental sampling, a single-year common garden and a multi-year reciprocal translocation between the *G. carneus* ecotypes, to test:

1. Are there differences in the soil niche between the seven *Gladiolus carneus* ecotypes? and do three focal populations of *G. carneus* occupy distinct soil niches?
2. Do *Gladiolus carneus* ecotypes have a fitness advantage on their native soil relative to other ecotypes within a common garden?
3. Do *Gladiolus carneus* ecotypes show evidence of local adaptation in a reciprocal translocation across an elevational gradient?

## MATERIALS AND METHODS

### Study Species

*Gladiolus carneus* (Iridaceae) is a cormous geophyte native to the CFR (Figure 1A-C). The species can be divided into seven morphologically distinct ecotypes, with each occupying distinct geographic range and abiotic niche (Khoury et al., in press). The CFR is characterised by a Mediterranean climate with winter rainfall and hot, dry summers (Manning and Goldblatt, 2012). *G. carneus* is winter growing, with individuals emerging from the dormant period in April or May and growing until they flower, from August to early January (Khoury et al., in press). All individuals go dormant in the late summer and only emerge at the beginning of the wet growing season.

**Figure 1.**
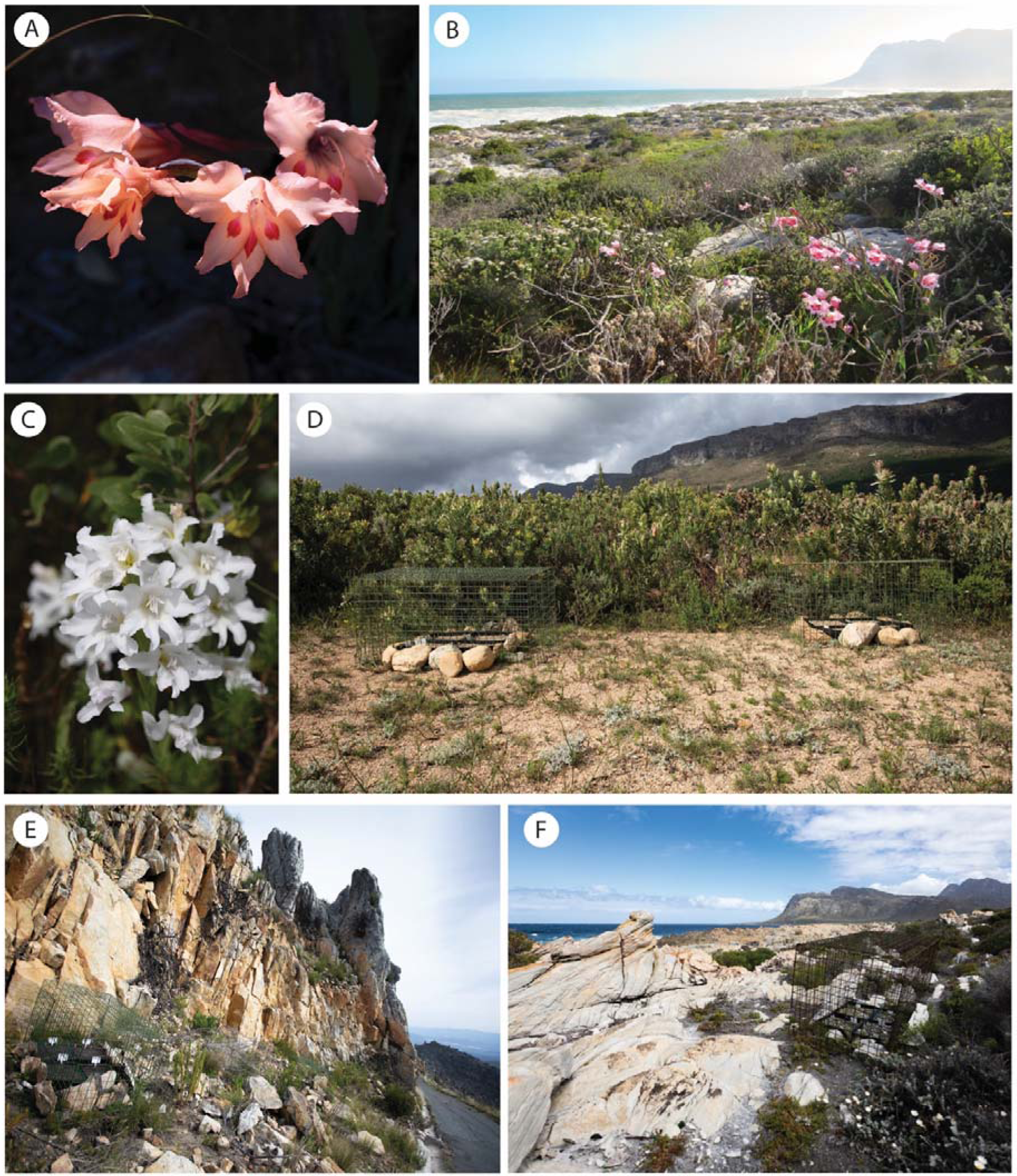
(A) Flowering *G. carneus* ecotypes from Jonaskop (JK), (B) Kleinmond Coast (KC), (C) and Limietberg (LB). Habitat images of reciprocal translocation sites (D) LB, (E) JK and (F) KC, with experimental plants in the foreground. Photos: A, C, D, E, F: Ethan Newman; B: Katharine Khoury.

To test for local adaptation between different ecotypes, we focused on three sites (Figure 1D-F), namely Jonaskop (JK, −33.963432, 19.510247), Kleinmond Coast (KC, −34.345452, 19.008462), and Limietberg (LB, −33.707683, 19.053684), which represent large populations of three ecotypes: *high-altitude*, *blandus*, and *albidus*, respectively. These three focal populations occupy divergent abiotic niches along an elevational gradient (Khoury et al., in press). JK, the *high-altitude* site, represents the highest elevational extreme (1403 m) while KC, the *blandus* site, represents the lowest elevation (9 m) in the *G. carneus* range. LB (457 m) represents a mid-elevation site.

### Do different ecotypes of *Gladiolus carneus* occupy distinct soil niches?

#### Rangewide ecotypic differences in soil niche

We tested whether there were differences in the soil niche of the different *G. carneus* ecotypes using point localities and soil layers from Cramer et al. (2019). The soil layers from Cramer et al. (2019) were modelled specifically for the Greater Cape Floristic Region (GCFR) using georeferenced soil data mined from the literature. When compared to SoilGrids layers, which predicts soil layers at a global scale, Cramer et al. (2019)’s regionally specific soil layers more accurately represent the heterogeneity of soil properties and make more reliable predictions of the vegetation types (E.g., fynbos, renosterveld) in the GCFR. The soil layers include electrical conductivity (mS/m), extractable potassium (cmol^+^/kg), extractable sodium (cmol^+^/kg), extractable phosphorus (mg/kg), pH, total carbon (%) and total nitrogen (%).

We used iNaturalist observations to compile a database of point localities of all ecotypes of *G. carneus.* The observations were classified into seven distinct ecotypes identified in Khoury et al. (in press). We excluded any observations that (1) did not have clear photos of the flowers, (2) observations of closely related taxa that had been misidentified as *G. carneus,* (3) observations that did not fit the descriptions of previously described ecotypes or were likely hybrids, (4) observations that occurred outside of the natural range of the species within the CFR (E.g., Australia), and (5) observations that were not research grade. All observations were filtered to include coordinates with an open geoprivacy and an accuracy under 100m. In addition to iNaturalist data, we provided additional point localities for poorly documented ecotypes based on field observations.

Overall, we included 789 localities in total, consisting of 209 *albidus*, 33 *blandus*, 67 *callistus*, 34 *high-altitude*, 37 *langeberg*, 379 *macowanianus* and 27 *prismatosiphon* point localities. These point localities were used to extract the soil properties from the seven soil layers in Cramer et al. (2019) using the ‘*raster*’ package (Hijmans, 2024).

#### Population level differences in soil niche

To test if there were differences between the *G. carneus* populations’ soil niche, we collected soil samples from three experimental sites, JK, KC, and LB which correspond to the populations used in the common garden and reciprocal translocation experiments. At each site, we collected five soil samples that were 200g each which were placed into Ziplock bags. All soil samples were taken within approximately one metre of a *G. carneus* individual.

Samples were collected between the 20^th^ and the 23^rd^ of June 2024, the middle of the growing season of *G. carneus* and were then sent to Elsenburg, Western Cape Department of Agriculture for soil analyses. For each soil sample, potassium (mg/kg), sodium (mg/kg), phosphorus (mg/kg), pH, carbon (%), and NH_4_ nitrogen (%) was measured.

#### Statistical Analyses

We used Principal Component Analyses (PCAs) to determine if the *G. carneus* ecotypes and populations cluster separately based on their soil niche. We implemented the PCAs using the *prcomp* function from the ‘*stats*’ package (R Core Team, 2021) and tested whether the ecotypes cluster separately using a permutation MANOVA from the package ‘*vegan*’ (Oksanen et al., 2020). Depending on the normality and homoscedasticity of the soil properties, we used a series of Kruskal-Wallis tests and one-way ANOVAs to test for differences between the ecotypes’ soil properties, both across the species range, and between the experimental populations. In each Kruskal-Wallis test or ANOVA, the soil variable was set as the response and ecotype or population as the fixed effect. Either a Dunn’s test, implemented using the ‘*dunn.test’* package (Dinno, 2024), or a pairwise t-test, implemented using the *pairwise.t.test* function in the ‘*stats*’ package (R Core Team, 2021), was used to determine the significance between ecotypes and populations. A Bonferroni correction was applied to all pairwise comparisons to reduce the likelihood of Type I errors.

### Do different ecotypes of *Gladiolus carneus* have a fitness advantage on their native soil relative to other ecotypes within a common garden?

#### Common garden experiment

To test whether the *G. carneus* ecotypes have a fitness advantage on their native soil relative to other ecotypes, we set up a common garden to track seedling survival and height (mm) in 2024, at the Department of Botany, Rhodes University. As the common garden used soil from each of the ecotypes’ native sites, we were testing whether there were fitness differences due to differences in the entire edaphic niche and not exclusively nutrient composition. In November and December 2023 and January 2024, we collected seeds from *G. carneus* populations from JK, KC, and LB. Each site was visited between four and six weeks after the peak flowering period where seeds were collected from dried-out, mature fruits of *G. carneus*. At JK, mesh organza bags were placed over flowering individuals that showed signs of early fruit development and were left to mature. Seeds were then collected roughly eight weeks after peak flowering from the bagged individuals. At each population, between 200 and 1 000 seeds were collected from 10 or more individuals. In the lab, the seeds were thoroughly mixed and stored in glass jars. On 1 May 2024, at the start of the next growing season of plants in the wild, seeds from the three populations were sowed in a 50:50 mix of pool sand and filtered soil collected from Mountain Drive, Makhanda (−33.328830, 26.507832) on which fynbos vegetation grows, the same vegetation type shared amongst the three populations. The growing medium was filtered using a stainless steel, mesh riddle with 0.5 x 0.5 cm squares to remove rocks, organic matter and large clumps of soil. This soil mixture resembles the topmost layer of soil in which the grit creates an airy medium where the seeds can germinate (Rachel Saunders, personal communication). This soil mixture was placed in seed trays (27 x 30 x 11.5 cm). Between 200 to 300 seeds were sprinkled on top of the soil and lightly covered in the soil-pool sand mix. Each tray contained seeds from only a single population. The trays were placed on a metal growing stand in a sunny courtyard of the Department of Botany where they were exposed to natural temperature fluctuations, often below 10°C in the evenings and above 20°C during the daytime, similar to the conditions in their native range. Seedlings experienced natural winter rainfall, as experienced in Makhanda. However, in periods without regular rainfall, the seeds were kept moist by watering the seedlings approximately every two days using stored rainwater. The seeds took approximately six weeks to germinate.

Between the 20^th^ and the 23^rd^ of June 2024, 20 litres of soil was collected from JK, KC and LB. Similarly, the soil from each site was filtered and placed into black seed trays. In addition to the trays containing soil from the *G. carneus* sites, we also filled separate trays with the medium in which the seeds were germinated to control for transplant shock. If the seedling did not experience transplant shock, survival rates after being transplanted from their germination medium into the control soil should be high. Any reduction in seedling survival in another soil medium would therefore be due to the differential soil properties and not due to being transplanted. In total, there were three trays of soil for each *G. carneus* site, and three trays of the control soil. On the 7^th^ of July 2024, we transplanted 30 germinated *G. carneus* seedlings from a single site into each tray. The 30 seedlings were evenly spaced out in six rows by five columns. Seedlings from all three *G. carneus* populations were transplanted into the four different soil types. All trays were placed on a single table a courtyard and rotated monthly. They were exposed to the same temperature and watering conditions as the germination trays.

Under greenhouse conditions with limited competition, frequent watering and nutrient-rich soil, *G. carneus* takes three years to flower (K. Khoury, unpublished data). Therefore, fitness measures such as the number of flowers or the number of seeds would only be used in long-term experiments on *G. carneus*. Instead, we used fitness components relevant to the early life stages, namely the seedling survival and height (*see* Wadgymar et al., 2024 for defining fitness through life history stages). These fitness measures were recorded once every four weeks for each individual in the common garden. Seedling height was measured from the soil to the tip of each leaf using digital calipers (0-200mm, TA). The last survival and height measurement for the year, taken on the 23^rd^ of September 2024, was used in all subsequent analyses as it represented survival and growth to the end of 2024. Measurements were not taken later in the year as *G. carneus* naturally dies back in late spring and summer.

#### Statistical Analyses

We used a series of generalized linear models (GLMs) to test if there were differences in fitness between the three ecotypes, JK, KC, and LB, on all soil types (*see Table S1 for sample sizes used in each analysis*). To test if there were differences in survival between the ecotypes, we used a binomial GLM with soil type, ecotype and their interaction as explanatory terms. In the model, survival was treated as a binary response variable (1 and 0). In the survival analysis, all four soil types were included (JK, KC, LB and the control soil) to test if there was any evidence of transplant shock. Thereafter, the control soil was excluded from analyses as there was no evidence of transplant shock (Figure 3A). To test if there were differences in the seedling’s height at the end of 2024, we modelled height using both gamma and Gaussian distributions. In both models, soil type, ecotype and their interaction were specified as explanatory variables. Any height measurements of seedlings that did not survive at the end of 2024 was excluded from the data set. The model diagnostics were checked using the *simulateResiduals* and *testDispersion* functions in the ‘*DHARMa*’ package (Hartig, 2022). The model with the gamma distribution was selected due to the lower AIC score. The *Anova* function from ‘*car*’ package (Fox and Weisberg, 2019) was used determine model significance for both the survival and height analysis. Furthermore, the *emmeans* function from the ‘*emmeans*’ package (Lenth, 2022) was used to conduct pairwise comparisons between the ecotypes on each soil type. A Bonferroni correction was applied to the pairwise comparisons to reduce the likelihood of Type 1 errors when making multiple comparisons.

For the survival analysis, we also used the ‘*emmeans*’ function to obtain the average probability of survival, plotted together with asymmetric confidence intervals.

In addition to investigating the effects of each fitness component using GLMs, we used aster models to test if there were differences in lifetime fitness. Aster models estimate lifetime fitness by taking into account multiple fitness measures, each with different distributions, that are dependent on the previous life stage (Geyer et al., 2007; Shaw et al., 2008). We used two fitness components in the aster model, survival at the end of 2024, modelled with a Bernoulli distribution (0 and 1), and height at the end of 2024, modelled using a normal distribution (Figure S1A). A normal distribution was used for height, as it is the only distribution available for continuous data within the aster framework. Any individuals that did not survive were given a height of 0 mm as the aster framework does not allow for missing data (Geyer et al., 2007; Shaw et al., 2008; Popovic and Lowry, 2020). The soil type, ecotype and their interaction were set as fixed factors within the aster models. The significance of fixed effects was determined by comparing a series of nested null models using likelihood ratio tests (Geyer et al., 2007; Shaw et al., 2008). The full aster model was used to predict the mean height of each ecotype on every soil type at the end of 2024, which takes into account survival. All analyses were conducted using the package, ‘*aster*’ (Geyer et al., 2007).

### Do different ecotypes of *Gladiolus carneus* show evidence of local adaptation across an elevational gradient?

#### Reciprocal translocation

To test whether the *G. carneus* ecotypes have a fitness advantage at their home site relative to other ecotypes, we set up a reciprocal translocation between Jonaskop (JK), Kleinmond Coast (KC), and Limietberg (LB). As the reciprocal translocation encompassed both the climatic and soil conditions for each site, we tested whether each of these ecotypes are locally adapted to their entire abiotic niche.

In November and December 2022 and January 2023, *G. carneus* seeds were collected from the three sites. The seeds were germinated in May 2023, at the Department of Botany, Rhodes University. The seed collection and germination followed the same procedure described for the common garden experiments. Between the 2^nd^ and 9^th^ of July 2023, a reciprocal translocation was set up between the three sites. From each of these sites, 120 litres of soil was collected in order to fill 16 or 20 black seed trays (27 x 30 x 11.5 cm) that were used in the reciprocal translocations. Soil from KC and JK was used fill 20 seed trays, while soil from LB filled 16 trays. Each tray contained eight seedlings from each locality which were transplanted into eight rows and four columns. The position of the seedlings from each locality was rotated through the trays. The trays were transported to each of the sites and set up under cages made of wire mesh that were designed to prevent large herbivores and rock falls from damaging the experiments. Cages were 1.2 x 0.6 x 0.8 m with 4 cm wide square holes painted with green rubberised paint to prevent rust. Each cage contained four randomly arranged trays, with each tray having a distinct arrangement of the seedlings. The cages were held in place by a combination of metal pegs and large rocks. Five cages were set up at the two most elevationally extreme sites, JK and KC, and four cages were set up at LB. Overall, 480 seedlings, consisting of 160 seedlings from each locality, were placed at JK and KC, while 384 seedlings with 128 from each locality were set up at LB. The cages were nested within the *G. carneus* populations at each site.

Similar to the common garden, seedling survival and height were documented as the fitness measures. Both survival and height were documented twice in 2023 and 2024, once during the middle of the growing season and once towards the end. Seedling survival was scored as a binary variable (0 or 1). The height (mm) of each seedling was measured in a straight-line distance from the surface of the soil to the tip of the leaf using digital callipers. Height measurements were not taken for individuals with evidence of damage on the leaf tips. The survival and height at the end of 2023 and 2024 were used in all analyses as this represented fitness at the end of each year.

#### Statistical Analyses

A series of generalized linear mixed models (GLMMs) were used to test whether there were differences in survival and height in both years between JK, KC and LB seedlings at each of their native sites. Survival in 2023 and 2024 were both modelled using a binomial error distribution while height (mm) in 2023 and 2024 were modelled using a normal distribution. All models had site, ecotype and their interaction as fixed factors. The models also included a nested random factor, tray within cage. Due to low sample sizes in the height measurements in 2024 (*see Table S2*) only cage, and not tray within cage, was included as a random factor in the 2024 height model. All models were implemented using the ‘*glmmTMB*’ package (Brooks et al., 2017). The model diagnostics were checked using the same procedure described for the common garden experiment. A type III ANOVA from the ‘*car*’ package (Fox and Weisberg, 2019) was used to determine the significance of fixed factors and pairwise contrasts between ecotypes at each site were conducted using the *emmeans* function from the ‘*emmeans*’ package (Lenth, 2022).

Similar to the common garden experiments, aster models were used to test for differences in the lifetime fitness of the seedlings at each reciprocal translocation site. We used three sets of aster models to look at differences in fitness in 2023, 2024 and both years in a full model. In the 2023 and 2024 models the fitness components included survival and height in their respective years, while the full model incorporated all four fitness components (Figure S1B). Survival was modelled using a Bernoulli distribution (0 or 1), while height was modelled using a normal distribution. Height in both years were set as terminal nodes within the models. In cases where seedlings did not survive, the height was set to 0 mm. Within the models, site, ecotype and their interaction were set as fixed effects. The significance of fixed effects was determined using likelihood ratio tests described under the common garden experiment. The three sets of aster models were used to make predictions about the height of the ecotypes at each site by the end of each year and while taking into account the previous dependencies. All data analyses was conducted in R v.4.1.2 (R Core Team, 2021) and the package ‘*ggplot2*’ (Wickham, 2016) was used for all data visualisation.

## RESULTS

### Do different ecotypes of *Gladiolus carneus* occupy distinct soil niches?

#### Rangewide ecotypic differences in soil niche

The permutation MANOVA showed that there were statistical differences between the *G. carneus* ecotypes’ soil niche (*F* = 33.35, *df* = 6, *P* < 0.001, Figure 2A). There were also statistical differences between the *G. carneus* ecotypes’ electrical conductivity (χ*2* = 117.75, *df* = 6, *P* < 0.0001, Table S3), extractable potassium (χ*2* = 298.80, *df* = 6, *P* < 0.0001, Table S3), extractable sodium (χ*2* = 198.94, *df* = 6, *P* < 0.0001, Table S3), extractable phosphorus (χ*2* = 281.98, *df* = 6, *P* < 0.0001, Table S3), pH (χ*2* = 147.28, *df* = 6, *P* < 0.0001, Table S3), total carbon (χ*2* = 189.06, *df* = 6, *P* < 0.0001, Table S3) and total nitrogen (χ*2* = 89.38, *df* = 6, *P* < 0.0001, Table S1). Pairwise comparisons showed there were significant differences between the ecotypes for all soil properties (*see Table S4*). These results suggest that there are differences in the ecotypes’ soil niche.

**Figure 2.**
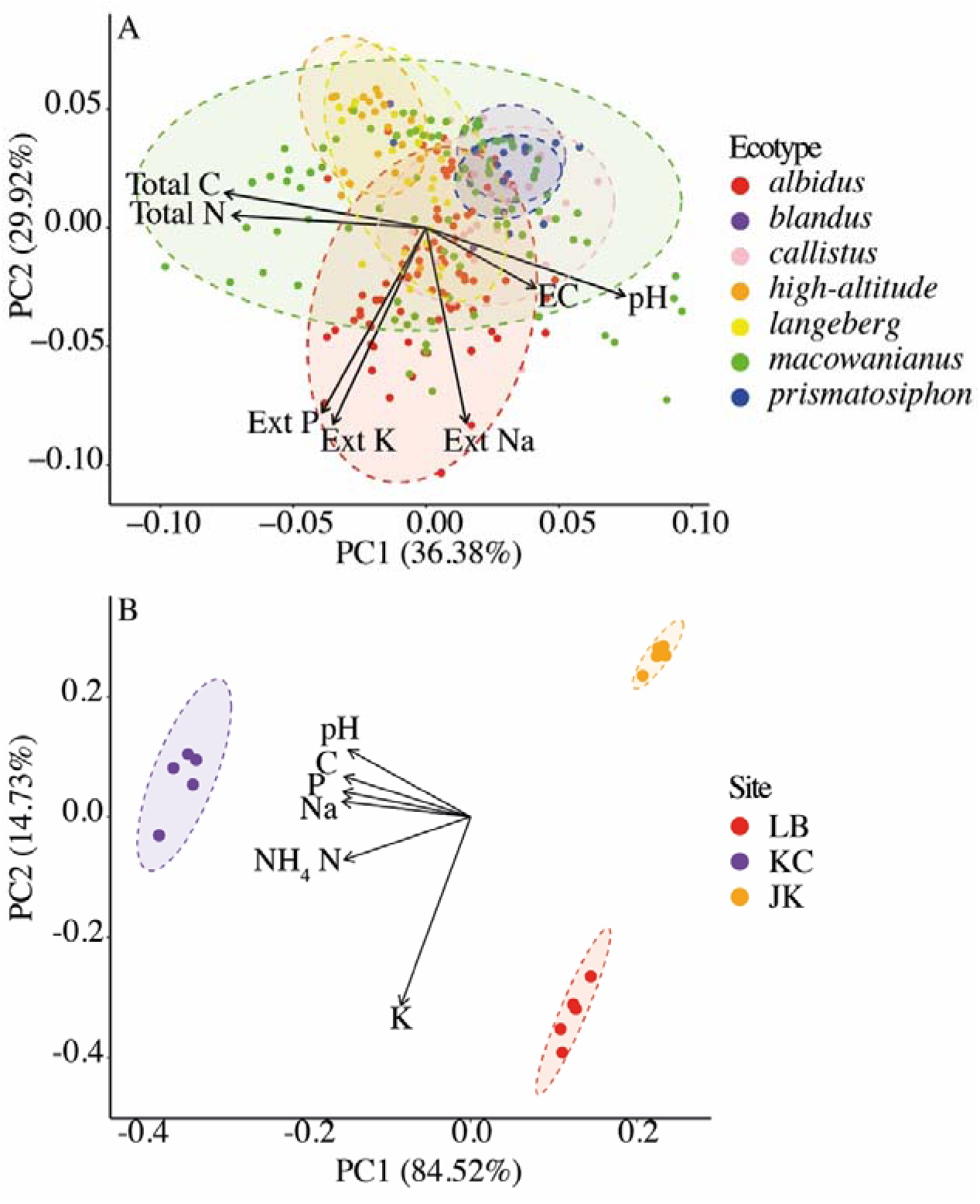
PCAs of the soil niches of the (A) *Gladiolus carneus* ecotypes, and (B) the three focal populations used in the common garden and reciprocal translocations. Both PCAs include 95% confidence ellipses and biplots of trait loadings. The variables included in the ecotypes’ soil niche includes, electrical conductivity (mS/m), extractable potassium (cmol^+^/kg), extractable sodium (cmol^+^/kg), extractable phosphorus (mg/kg), pH, total carbon (%), and total nitrogen (%) extracted from Cramer et al. (2019). The variables used to test for differences between the populations’ soil niche includes, potassium (mg/kg), sodium (mg/kg), phosphorus (mg/kg), pH, carbon (%), and NH_4_ Nitrogen (%).

#### Population level differences in soil niche

The permutation MANOVA showed that there were statistical differences between the soil properties at JK, KC, and LB (*F* = 494.90, *df* = 2, *P* < 0.001, Figure 2B). In particular, there were differences in the potassium (*F* = 251.70, *df* = 2, *P* = 0.0001, Table S5), sodium (χ*2* = 12.59, *df* = 2, *P* = 0.0018, Table S5), phosphorus (χ*2* = 12.39, *df* = 2, *P* = 0.0020, Table S5), pH (χ*2* = 458.60, *df* = 2, *P* < 0.0001, Table S5), carbon (χ*2* = 10.92, *df* = 2, *P* = 0.0043, Table S5), and NH_4_ nitrogen (*F* = 603.50, *df* = 2, *P* < 0.0001, Table S5) measured at those three sites. Pairwise comparisons further showed there were significant differences between the sites for all soil properties documented (*see Table S6*). KC had the highest sodium (283.60 ± 9.68 SE mg/kg), phosphorus (107.60 ± 5.07 SE mg/kg), carbon (7.19 ± 0.21 SE %), NH_4_ nitrogen (0.41 ± 0.01 SE %) and pH (5.52 ± 0.04 SE) indicating that it is a ‘nutrient-rich’ site (Table S5). JK had the lowest potassium (28.00 ± 1.34 SE mg/kg), sodium (9.00 ± 0.63 SE mg/kg), phosphorus (13.60 ± 0.68 SE mg/kg) and NH_4_ nitrogen (0.03 ± 0.01 SE %) indicating that it is a ‘nutrient-poor’ site (Table S5). Additionally, LB had the highest potassium (95.20 ± 2.35 SE mg/kg) but lowest pH (4.50 ± 0.00SE) and carbon (0.62 ± 0.01 SE %) but was otherwise considered as a site with intermediate nutrients (Table S5). Overall, there are differences between the soil properties of the three sites, which represent a gradient between nutrient-rich and poor sites within the *G. carneus* species complex.

### Do different ecotypes of *Gladiolus carneus* have a fitness advantage on their native soil relative to other ecotypes within a common garden?

#### Common garden experiment

Overall, all seedlings in the common garden did not have a survival advantage, but did have a growth advantage on nutrient-rich KC compared to the other two soils (Figure 3). Both KC and JK seedlings, which occupy the most extreme soil niches, had a fitness advantage on their native soils, while LB seedlings did not have fitness advantage on their native soil compared to JK and KC seedlings (Figure 3C).

Seedlings experienced high survival rates across all ecotypes on all soil types at the end of 2024 (Figure 3A). This included seedlings on the control soil indicating that there was no evidence of transplant shock (Figure 3A). Soil type (χ*2* = 13.30, *df* = 3, *P* = 0.0040, Figure 3A) and ecotype (χ*2* = 12.13, *df* = 2, *P* = 0.0023, Figure 3A) were significant predictors of survival in 2024, whilst the interaction between soil type and ecotype was not (χ*2* = 8.04, *df* = 6, *P* = 0.2355, Figure 3A). The pairwise comparisons between the ecotypes on each soil type were all non-significant (*P* = 1.00, Table S7), indicating that there were no differences in survival between seedlings.

**Figure 3.**
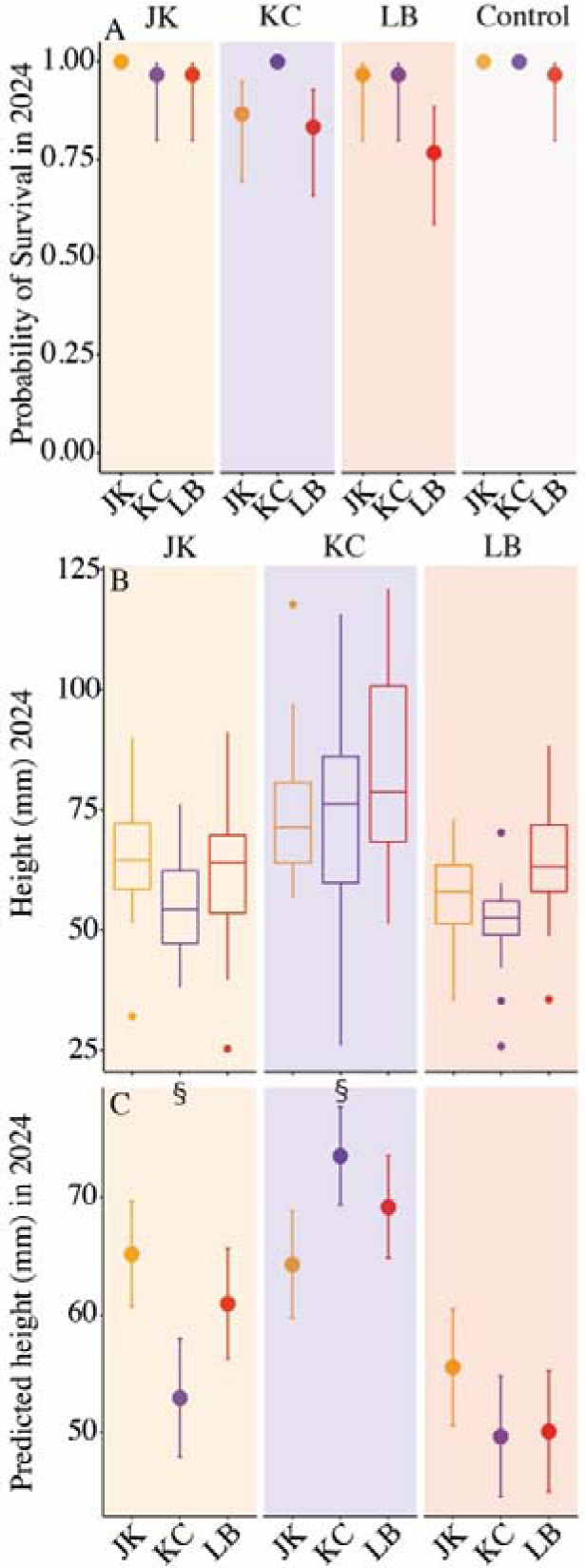
Common garden: (A) The probably of survival, (B) height (mm), and (C) the predicted height (mm) at the end of 2024 of the seedlings in the common garden. Jonaskop (JK), Kleinmond Coast (KC), and Limietberg (LB) seedlings were transplanted into JK, KC, LB, and control soil. The soil types are represented by the bands through the figure. In A, the round points represent the predicted probably of survival with asymmetric confidence intervals. In C, the round points represent the mean height and standard error as predicted by the aster analysis. The ‘§’ indicates the native ecotype has a fitness advantage based on foreign vs local criteria for local adaptation.

Soil type (χ*2* = 94.22, *df* = 2, *P* < 0.0001, Figure 3B), ecotype (χ*2* = 25.72, *df* = 2, *P* < 0.0001, Figure 3B) and their interaction (χ*2* = 10.88, *df* = 4, *P* = 0.03, Figure 3B) were all significant predictors of seedling height at the end of 2024. The pairwise comparisons showed that JK seedlings were significantly larger than the KC seedlings on JK soil (*P* < 0.05, Table S7) and LB seedlings were significantly larger than KC seedlings on LB soil (*P* = 0.0007, Table S7). These results indicate that JK and LB seedlings have an advantage over KC seedlings on their native soil. There were no other significant differences between the seedling heights.

The aster model showed that soil type (*P* < 0.0001, Table 1, Figure 3C) was a significant predictor of lifetime fitness, but ecotype (*P* = 0.75, Table 1, Figure 3C) and the interaction between soil type and ecotype (*P* = 0.21, Table 1, Figure 3C) were not. Seedlings from JK and KC had the highest predicted mean height on their native soil, however, LB seedlings did not show the same homesite fitness advantage (Figure 3C).

**Table 1:**
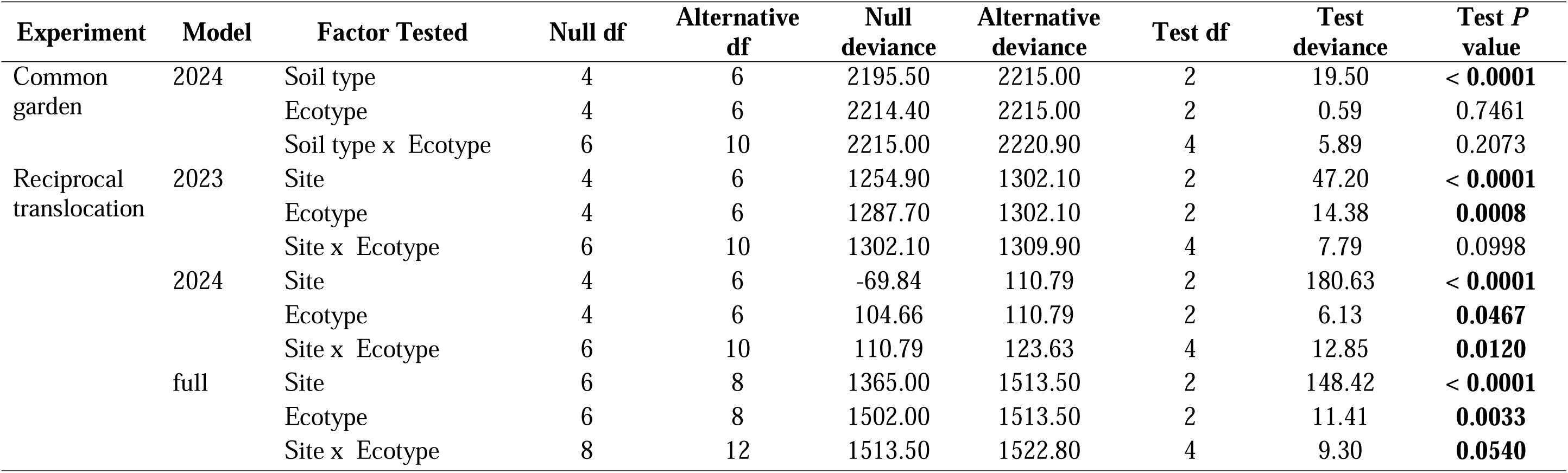
Aster model results from the common garden and reciprocal translocation experiments. The common garden experiment includes survival and height (mm) in 2024. Three aster model results are presented for the reciprocal translocation, the first includes the survival and height (mm) in 2023, the second includes survival and height (mm) in 2024, and the third includes data from both years.

| Experiment | Model | Factor Tested | Null df | Alternative df | Null deviance | Alternative deviance | Test df | Test deviance | Test *P* value |
| --- | --- | --- | --- | --- | --- | --- | --- | --- | --- |
| Common garden | 2024 | Soil type | 4 | 6 | 2195.50 | 2215.00 | 2 | 19.50 | < **0.0001** |
|  |  | Ecotype | 4 | 6 | 2214.40 | 2215.00 | 2 | 0.59 | 0.7461 |
|  |  | Soil type x Ecotype | 6 | 10 | 2215.00 | 2220.90 | 4 | 5.89 | 0.2073 |
| Reciprocal translocation | 2023 | Site | 4 | 6 | 1254.90 | 1302.10 | 2 | 47.20 | < **0.0001** |
|  |  | Ecotype | 4 | 6 | 1287.70 | 1302.10 | 2 | 14.38 | **0.0008** |
|  |  | Site x Ecotype | 6 | 10 | 1302.10 | 1309.90 | 4 | 7.79 | 0.0998 |
|  | 2024 | Site | 4 | 6 | -69.84 | 110.79 | 2 | 180.63 | < **0.0001** |
|  |  | Ecotype | 4 | 6 | 104.66 | 110.79 | 2 | 6.13 | **0.0467** |
|  |  | Site x Ecotype | 6 | 10 | 110.79 | 123.63 | 4 | 12.85 | **0.0120** |
|  | full | Site | 6 | 8 | 1365.00 | 1513.50 | 2 | 148.42 | < **0.0001** |
|  |  | Ecotype | 6 | 8 | 1502.00 | 1513.50 | 2 | 11.41 | **0.0033** |
|  |  | Site x Ecotype | 8 | 12 | 1513.50 | 1522.80 | 4 | 9.30 | **0.0540** |

### Do different ecotypes of *Gladiolus carneus* show evidence of local adaptation across an elevational gradient?

#### Reciprocal Translocation

Over the two years, three cages were removed from the data set (*see resulting sample sizes in Table S2*). At the KC site, two of the cages were placed approximately 50m from the shoreline representing the edge of the native population, and nearly all the seedlings in those cages died within the first year. A third cage showed signs of human tampering. All three cages were excluded from analyses. Additionally, any seedlings showing signs of damage were excluded.

There were relatively high survival rates (>70%) of all ecotypes at every site in 2023, however, this was not the case in 2024 (Table S8). In 2023, site (χ*2* = 11.97, *df* = 2, *P* = 0.0025, Figure 4A), ecotype (χ*2* = 19.31, *df* = 2, *P* < 0.0001, Figure 4A) and their interaction (χ*2* = 10.53, *df* = 4, *P* = 0.0324, Figure 4A) were all significant predictors of seedling survival. Similarly, in 2024, site (χ*2* = 19.46, *df* = 2, *P* < 0.0001, Figure 4B), ecotype (χ*2* = 16.90, *df* = 2, *P* = 0.0002, Figure 4B) and their interaction (χ*2* = 12.64, *df* = 4, *P* = 0.0132, Figure 4B) were all significant predictors of seedling survival. At JK in 2023 and 2024, the native JK seedlings had significantly higher probability of survival than KC seedlings (P < 0.01, Table S9). Additionally, JK, KC and LB seedlings all had the highest survival rates at their native sites in 2024 (Figure 4B, Table S8).

**Figure 4.**
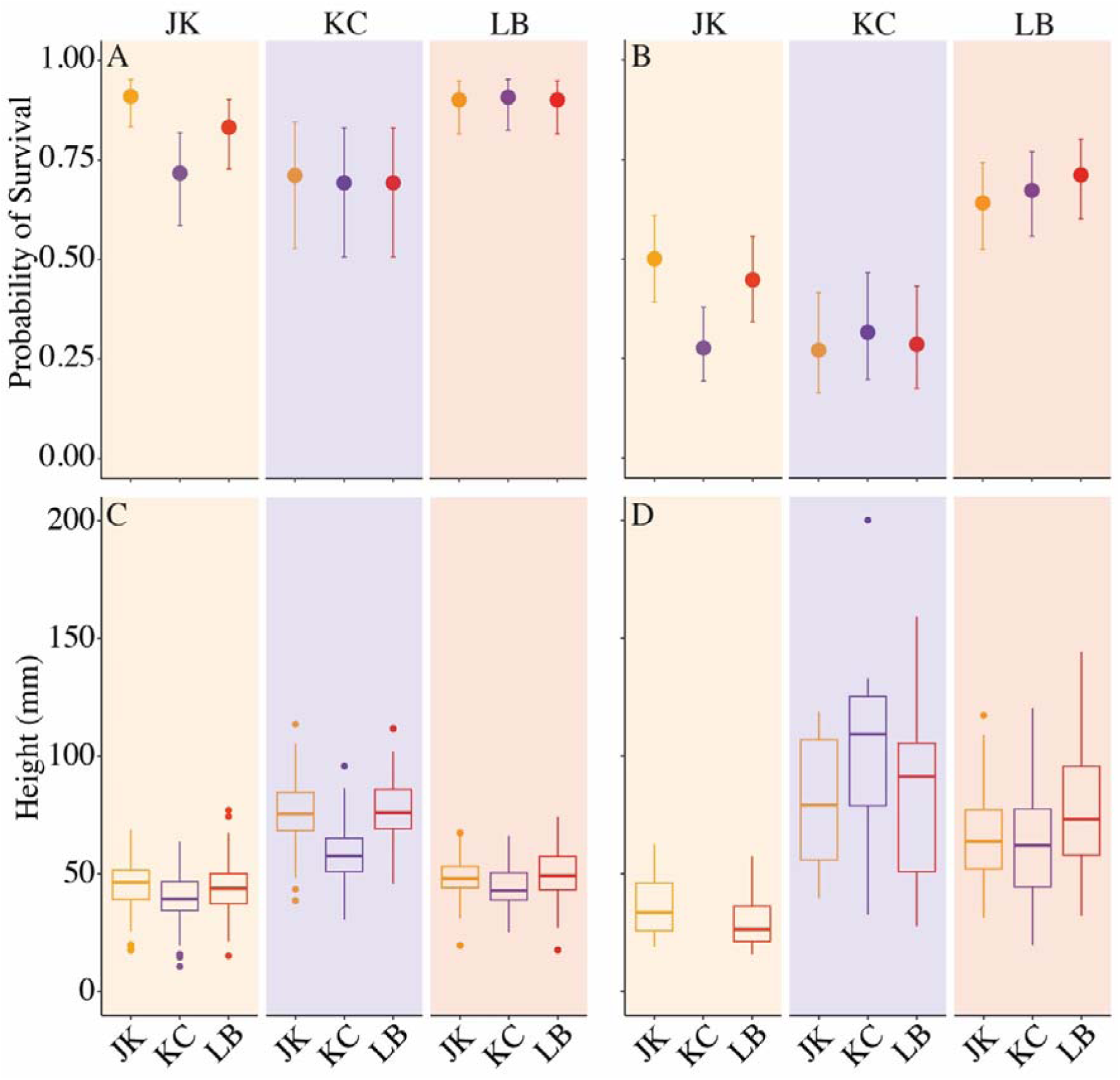
Reciprocal translocation: The probably of survival in (A) 2023 and (B) 2024, and height (mm) at the end of (C) 2023 and (D) 2024 between the three reciprocal translocation sites, Jonaskop (JK), Kleinmond Coast (KC), and Limietberg (LB). Generalized linear mixed models were used to model the effects of site, ecotype and their interaction on survival and height (mm) in 2023 and 2024. The sites are represented by the bands through the figure. The circles in A and B represent the predicted survival with asymmetric error bars.

In 2023, site (χ*2* = 197.68, *df* = 2, *P* < 0.0001, Figure 4C), ecotype (χ*2* = 20.42, *df* = 2, *P* < 0.0001, Figure 4C) and their interaction (χ*2* = 28.75, *df* = 4, *P* < 0.0001, Figure 4C) were all significant predictors of seedling height. JK and LB seedlings were significantly larger than KC at JK (P < 0.05, Table S9). LB seedlings were significantly larger than KC at LB (P < 0.05, Table S9). Both JK and LB seedlings were significantly larger than KC at KC (P < 0.0001, Table S9). These results indicate that JK and LB seedlings have a fitness advantage over KC at their native sites. Additionally, JK and LB seedlings were both significantly larger than KC seedlings at KC, indicating that two introduced localities had a fitness advantage over the native (P < 0.0001, Table S9).

In 2024, site (χ*2* = 23.03, *df* = 2, *P* < 0.0001, Figure 4D) and the interaction between site and ecotype (χ*2* = 20.38, *df* = 4, *P* < 0.0004, Figure 4D) were significant predictors of height, however, ecotype was non-significant (χ*2* = 0.08, *df* = 2, *P* = 0.9594, Figure 4D). LB seedlings were significantly larger than KC seedlings at LB (P < 0.05, Table S9). There were no other comparisons of the ecotypes’ heights that were significantly different. Overall, native seedlings were, on average, the largest seedlings at their native site in 2024 (Figure 4D).

We used aster models to test whether there were differences in the lifetime fitness of the seedlings and to make predictions about the seedlings’ heights at each reciprocal translocation site. The first and second aster models took into account the survival and height data from 2023 and 2024, respectively. These models were then used to make predictions of the seedling heights using only data from those years. The third full aster model took into account the seedling survival and height from 2023 and 2024, and the associated height predicted for the end of 2024 takes into account both years’ data. The 2023 aster model showed that site (*P* < 0.0001) and ecotype (*P* < 0.001), but not their interaction (*P* = 0.0998), were significant predictors of lifetime fitness. Furthermore, only JK seedlings had a fitness advantage at their native site in 2023. The 2024 and the full aster models showed that site (2024: *P* < 0.0001, full: *P* < 0.0001), ecotype (2024: *P* < 0.05, full: *P* < 0.01) and their interaction (2024: *P* < 0.05, full: *P* = 0.05) were all significant predictors of lifetime fitness (Table 1, Figure 5). In the 2024 model, all native seedlings were predicted to have a home-site advantage, while the full model suggested that JK and LB plants had a fitness advantage in their native environment while KC plants did not have a fitness advantage at their native site (Figure 5C). These results suggest that there is evidence for local adaptation at all three sites, with the strongest evidence for local adaptation being at the JK, the nutrient-poor site, and LB, the intermediate nutrient site.

**Figure 5.**
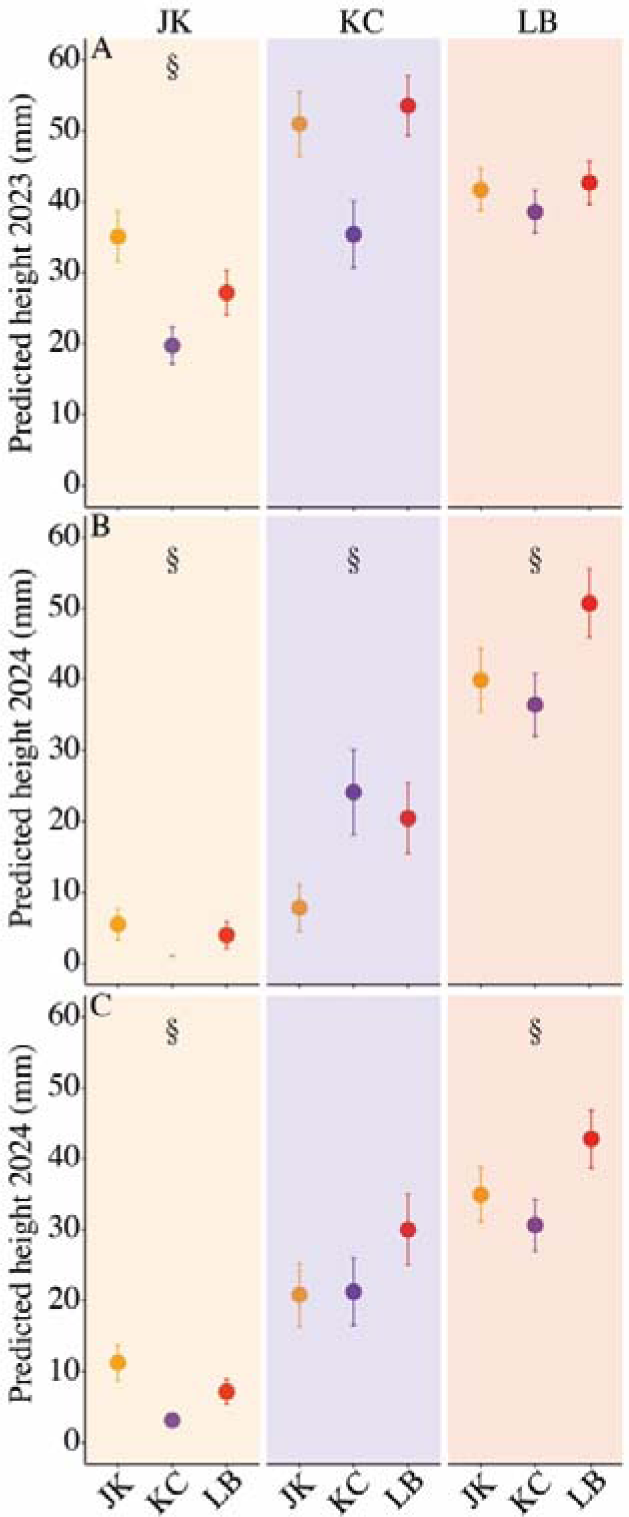
Reciprocal translocation: Predicted mean height (mm) of the (A) 2023 aster model, (B) the 2024 aster model and (C) the full 2023 and 2024 model. The 2023 and 2024 models only take into account the survival and height (mm) within their respective years. The full model takes into account survival and height in both years. All three models show the predicted height for ecotypes at the three reciprocal translocation sites, Jonaskop (JK), Kleinmond Coast (KC), and Limietberg (LM). The sites are represented by the bands through the figure. The circles represent the expected mean height with standard error. The ‘§’ indicates the native ecotype has a fitness advantage based on foreign vs local criteria for local adaptation.

## DISCUSSION

This study provides evidence that the *G. carneus* ecotypes occupy distinct soil niches and that they are in part, locally adapted to these soil niches at the seedling stage. This is the first study providing evidence for local adaptation of plants occupying different soil niches in the CFR using both a common garden experiment that isolates the effects of soil properties from climatic variables on seedling survival and height, and a reciprocal translocation to test for local adaptation. Interestingly, and in agreement with the broader literature (*see* Sambatti and Rice, 2006; Lekberg et al., 2012; Ferris and Willis, 2018), we found local adaptation between localities with extreme soil properties, namely between extremely nutrient-poor soils and nutrient-rich soils. As predicted, all ecotypes had a growth advantage on the on the nutrient-rich soils. However, the reciprocal translocation also revealed evidence of local adaptation at a locality with intermediate soil nutrient composition. However, in this case, it is likely that other abiotic factors, rather than soil type, is contributing to local adaptation, as the results from the reciprocal translocation experiment were not entirely supported by the common garden experiment. Below we discuss the results in the context of the broader literature with the implications for the diversification of the Cape Floristic Region, a biodiversity hotspot.

### Do *Gladiolus carneus* ecotypes occupy distinct soil niches?

All seven of the *G. carneus* ecotypes, and the three experimental populations, all occupied distinct soil niches (Figure 2). The three experimental populations occupied a gradient of soil niches, which included a nutrient-poor site (JK), one intermediate site (LB) and one nutrient-rich site (KC) (Figure 2B). These results provide evidence that shifts in soil nutrient composition between both ecotypes and populations are common in the species complex and is potentially a driver of diversification. Similar shifts have been documented between populations of the same species, occupying distinct soil niches (Sambatti and Rice, 2006; Jimenez-Ambriz et al., 2007; Macel et al., 2007; Lekberg et al., 2012; Dittmar and Schemske, 2023; Jimenez-Ramirez et al., 2023) or between closely related species (Verboom et al., 2004; Ellis et al., 2006) and even between populations within a wider species complex (Verboom et al., 2012). Within the CFR, shifts in soil type between sister species have commonly been shown using molecular phylogenies and frequently show that edaphic shifts between sister taxa are common (van der Niet and Johnson, 2009; Schnitzler et al., 2011; Forest et al., 2014). These shifts in soil type are then inferred as a major driver of speciation.

### Are *Gladiolus carneus* ecotypes locally adapted to their soil niches?

Despite all the populations at the translocation sites occupying distinct edaphic niches, the common garden experiment indicated that only JK and KC, which occupied the extreme soil niches, showed any evidence of local adaptation to their edaphic niche. In contrast, LB, which occupied an intermediate edaphic niche, showed no evidence local adaptation to its edaphic niche. Similarly, Jimenez-Ramirez et al. (2023) found no evidence of local adaptation to non-extreme soils over small spatial scales during the early life stages of *Pinus sylvestus*. These results indicate that shifts in soil niche, and particularly non-extreme soil shifts, do not necessarily result in local adaptation. Furthermore, edaphic shifts often co-occur with climatic shifts, which may result in local adaptation to exclusively to climate (Macel et al., 2007; Ellis and Ågren, 2024). As soil shifts do not necessarily result in local adaptation, they should be used cautiously when inferring speciation.

Additionally, the two populations showing evidence of local adaptation to their soil niche (JK and KC) also occupy the two elevational extremes of the *G. carneus* range and are thus exposed to other abiotic factors that may be contributing to local adaptation. In particular, the JK population is exposed to freezing temperatures and snow during the winter growing season, and the KC population grows along the coast and is frequently exposed to salt spray. Both freezing temperatures (Agren and Schemske, 2012) and salt spray (Lowry et al., 2008; Busoms et al., 2015; Popovic and Lowry, 2020) have been shown to exert strong selection that can result in local adaptation, which may be contributing to the low survival rates at the JK and KC sites. The relative contribution of these ecological factors can be disentangled using factorial reciprocal translocations (*see* Macel et al., 2007; Popovic and Lowry, 2020; Ellis and Ågren, 2024). For example, Ellis and Ågren (2024) reciprocally transplanted populations of *Arabidopsis thaliana* and their soils between Italy and Sweden to isolate the contribution of soil type verses climate to local adaptation. Using this method, they found soil type made little contribution to local adaptation. Similar factorial reciprocal translocations could be employed between *G. carneus* populations to isolate the contribution of specific ecological factors to local adaptation.

This approach would be particularly useful for the LB population, which showed evidence of local adaptation to factors other than its edaphic niche. Other ecotypes within *G. carneus* may be locally adapted to other ecological factors that have not been explicitly tested (E.g.: climate, fire-regime, rocky outcrops), or to an interaction between their edaphic niche and other ecological factors. Dittmar and Schemske (2023) demonstrated such an interaction when they conducted a multi-year reciprocal translocation between *Leptosiphon parviflorus* populations on sandstone and serpentine soil. They found that the population native to the serpentine soil consistently had a home-site advantage, while the population native to the sandstone soil only had a home-site advantage in two of the four years. Using a greenhouse experiment, they further demonstrated that the sandstone population’s fitness advantage was dependent on water availability. Dittmar and Schemske (2023)’s results also highlight that the interactions causing local adaptation may vary temporally. Furthermore, the other *G. carneus* ecotypes may be locally adapted at a different life stage. Both the common garden and reciprocal translocations presented here only measured local adaptation from the seedling stage. However, early life stages (E.g.: germination) has been shown to strongly contribute to local adaptation (Postma and Agren, 2016). This underscores that multi-year reciprocal translocations, that include all life stages, are needed to understand the variability of selection over time and the extent of local adaptation (Wadgymar et al., 2022).

### Are edaphic shifts a driver of diversification in the Cape Floristic Region?

We have presented data on the first reciprocal translocation and common garden experiments explicitly testing for local adaptation to divergent soil types between closely related taxa in the CFR. Our results suggest that shifts in edaphic niches occur in the CFR and can result in local adaptation, especially in some extreme niches. The frequency of these edaphic shifts in the CFR is somewhat disputed with Schnitzler et al. (2011) showing edaphic shifts between 20 – 72% of sister species pairs from four Cape clades, and van der Niet and Johnson (2009) showing edaphic shifts in only 17% of sister species pairs from eight Cape clades.

Furthermore, shifts in substrate type have also been documented in *Lapeirousia* (Forest et al., 2014) and *Ehrharta* (Verboom et al., 2004). Despite these shifts being documented on a macroevolutionary scale, there is limited experimental evidence demonstrating shifts and there are no examples of reciprocal translocations or common garden experiments within the CFR explicitly testing for local adaptation between contrasting edaphic niches. The strongest experimental evidence comes from Verboom et al. (2004) which showed that seedlings from eight *Ehrharta* species grown under controlled nutrient-rich conditions, differed in their relative growth rates and were associated both with different adult stage growth forms (slow, intermediate and fast growers) and with their native edaphic conditions. Verboom et al. (2004) further showed that the shifts from the slow to the fast growth form only occurred after a transition from nutrient-poor, sandstone derived soils to richer, shale and granite derived soils. Furthermore, the only reciprocal translocation testing for local adaptation along an ecological gradient in the CFR, Latimer et al. (2009), did not isolate the edaphic niche from other abiotic factors. The only example of a reciprocal translocation testing for local adaptation between contrasting edaphic environments in the GCFR is that of Ellis and Weis (2006). They conducted a reciprocal translocation between *Argyroderma* species occupying distinct soil microenvironments in the Succulent Karoo and found evidence for local adaptation. Overall, these results are the first in the CFR showing both frequent edaphic shifts and finding evidence for local adaptation in contrasting edaphic niches.

## CONCLUSION

In this study, we present a single year of a common garden experiments testing for differential fitness on contrasting soil types, and two years of a reciprocal translocation testing for local adaptation. We found that only two of the three populations, JK and KC were locally adapted to their soil niche, while the LB was locally adapted its abiotic niche. Further research should identify ecological shifts and use multi-year common garden and reciprocal translocations, that include all life stages, to comprehensively test for local adaptation (Wadgymar et al., 2017). We should additionally be using factorial reciprocal translocations and common garden experiments to isolate the role of specific, or interacting abiotic (E.g. climatic, topographic, and edaphic) or biotic (E.g. pollinators and herbivores) factors driving causing local adaptation (Dittmar and Schemske, 2023; Ellis and Ågren, 2024). Furthermore, when ecological factors causing local adaptation are identified in the CFR, their contribution to pre and postzygotic reproductive isolation should be quantified to better understand their role in driving speciation in this hyper-diverse region (Baack et al., 2015; Rajakaruna, 2017).

## ACKNOWLEDGEMENTS

We thank Marelise Faul, Kaelin du Plessis, Jordan McCullough, and Tafadzwa Thabethe for field assistance. Robyn Khoury, Jamie Venter, Tafadzwa Thabethe, the Newman family and the Hansford family for providing accommodation. We thank Cape Nature (CN35-87-18949) for providing permits to conduct this research. The research was financially supported by the Botanical Education Trust (KLK) and NRF-Thuthuka Grant TTK210211585733 (ELN).

## AUTHOR CONTRIBUTIONS

KLK and ELN conceptualised the study. KLK collected and analysed the data. KLK wrote the paper with input from ELN.

## DATA AVALIABILITY STATEMENT

All data and code associated with this manuscript will be made publicly available upon acceptance.

## SUPPORTING INFORMATION

**Table S1**. Sample sizes for common garden experiments testing for differential fitness between Jonaskop (JK), Kleinmond Coast (KC), and Limietberg (LB) seedlings on each of their respective soils and a control soil.

**Table S2**. Sample sizes for the reciprocal translocation experiments testing for differential fitness between Jonaskop (JK), Kleinmond Coast (KC), and Limietberg (LB) seedlings at each of their respective native sites.

**Table S3.** The mean and standard error of soil layers from Cramer *et al*. (2019) for *Gladiolus carneus* ecotypes.

**Table S4.** Pairwise comparisons between the *Gladiolus carneus* ecotypes soil properties.

**Table S5.** The mean and standard error of soil measurements taken at Jonaskop (JK), Kleinmond coast (KC), and Limietberg (LB).

**Table S6.** Pairwise comparisons between the soil measurements at Jonaskop (JK), Kleinmond coast (KC), and Limietberg (LB).

**Table S7**. Pairwise comparisons for the survival and height (mm) of Jonaskop (JK), Kleinmond Coast (KC), and Limietberg (LB) seedlings on each soil type. Soil type includes the native soil of all respective plant localities and a control soil.

**Table S8.** The survival rate (%) of all ecotypes at all four translocation sites in 2023 and 2024. All survival rates presented are the percentage of seedlings that survived from the beginning of the experiment.

**Table S9**. Pairwise comparisons between Jonaskop (JK), Kleinmond Coast (KC), and Limietberg (LB) seedlings at all the reciprocal translocation sites. Generalized linear mixed models were used to test for differences in survival and height at the end of 2023 and 2024.

**Figure S1.** Aster model dependency structure used in the (A) common garden and (B) full 2023 and 2024 reciprocal translocation.

