## Supplements for "Local adaptation of *Gladiolus carneus* to soil nutrient extremes in the Cape Floristic Region"

### Supplementary materials

**Table S1**. Sample sizes for common garden experiments testing for differential fitness between Jonaskop (JK), Kleinmond Coast (KC), and Limietberg (LB) seedlings on each of their respective soils and a control soil.

| | | GLMM | | aster |
|---|---|---|---|---|
| **Soil type** | **Ecotype** | **Survival 2024** | **Height 2024 (mm)** | |
| JK | JK | 30 | 30 | 30 |
| JK | KC | 30 | 29 | 30 |
| JK | LB | 30 | 29 | 30 |
| KC | JK | 30 | 26 | 30 |
| KC | KC | 30 | 30 | 30 |
| KC | LB | 30 | 25 | 30 |
| LB | JK | 30 | 29 | 30 |
| LB | KC | 30 | 29 | 30 |
| LB | LB | 30 | 23 | 30 |
| Control | JK | 30 | - | - |
| Control | KC | 30 | - | - |
| Control | LB | 30 | - | - |

**Table S2**. Sample sizes for the reciprocal translocation experiments testing for differential fitness between Jonaskop (JK), Kleinmond Coast (KC), and Limietberg (LB) seedlings at each of their respective native sites.

| | | **GLMM** | | | **aster** |
|---|---|---|---|---|---|
| **Site** | **Ecotype** | **Survival 2023 & 2024** | **Height 2023 (mm)** | **Height 2024 (mm)** | |
| JK | JK | 160 | 119 | 12 | 75 |
| JK | KC | 160 | 88 | 0 | 99 |
| JK | LB | 160 | 96 | 15 | 82 |
| KC | JK | 64 | 36 | 6 | 47 |
| KC | KC | 64 | 33 | 12 | 43 |
| KC | LB | 64 | 44 | 13 | 54 |
| LB | JK | 128 | 100 | 67 | 110 |
| LB | KC | 128 | 96 | 73 | 106 |
| LB | LB | 128 | 94 | 77 | 108 |

**Table S3.** The mean and standard error of soil layers from Cramer *et al*. (2019) for *Gladiolus carneus* ecotypes.

| **Ecotype** | **Electrical conductivity (mS/m)** | **Ext Potassium (cmol⁺/kg)** | **Ext Sodium (cmol⁺/kg)** | **Ext Phosphorus (mg/kg)** | **pH** | **Total Carbon (%)** | **Total Nitrogen (%)** |
|---|---|---|---|---|---|---|---|
| *albidus* | 11.93 ± 7.51 | 0.45 ± 0.18 | 0.28 ± 0.14 | 13.51 ± 5.61 | 4.57 ± 0.49 | 2.07 ± 0.80 | 0.18 ± 0.06 |
| *blandus* | 14.28 ± 8.55 | 0.09 ± 0.03 | 0.1 ± 0.06 | 3.48 ± 2.44 | 5.19 ± 0.55 | 1.18 ± 0.93 | 0.19 ± 0.04 |
| *callistus* | 22.67 ± 19.12 | 0.13 ± 0.09 | 0.23 ± 0.11 | 5.56 ± 2.82 | 5.02 ± 0.78 | 1.77 ± 0.80 | 0.19 ± 0.04 |
| *high-altitude* | 8.42 ± 1.78 | 0.1 ± 0.08 | 0.08 ± 0.04 | 4.74 ± 3.76 | 3.71 ± 0.31 | 4.76 ± 1.23 | 0.32 ± 0.11 |
| *langeberg* | 6.82 ± 1.52 | 0.18 ± 0.10 | 0.14 ± 0.07 | 5.80 ± 3.40 | 3.75 ± 0.32 | 2.25 ± 1.63 | 0.24 ± 0.10 |
| *macowanianus* | 17.47 ± 20.32 | 0.18 ± 0.14 | 0.18 ± 0.07 | 7.52 ± 5.38 | 4.44 ± 1.00 | 3.52 ± 2.13 | 0.35 ± 0.28 |
| *prismatosiphon* | 7.59 ± 2.36 | 0.14 ± 0.06 | 0.19 ± 0.07 | 0.50 ± 0.39 | 5.04 ± 0.85 | 1.14 ± 0.86 | 0.12 ± 0.06 |

**Table S4.** Pairwise comparisons between the *Gladiolus carneus* ecotypes soil properties. Significant pairings are highlighted in bold.

| **Comparison** | **EC (mS/m)** | **Ext Potassium (cmol+/kg)** | **Ext Sodium (cmol+/kg)** | **Ext Phosphorus (mg/kg)** | **pH** | **Total Carbon (%)** | **Total Nitrogen (%)** |
|---|---|---|---|---|---|---|---|
| *albidus – blandus* | 0.1061 | **<0.0001** | **<0.0001** | **<0.0001** | **0.0073** | **<0.0001** | 1.0000 |
| *albidus – callistus* | **0.0001** | **<0.0001** | **<0.0001** | **<0.0001** | **0.0326** | 0.3930 | 1.0000 |
| *albidus – high-altitude* | 0.0870 | **<0.0001** | **<0.0001** | **<0.0001** | **<0.0001** | **<0.0001** | **<0.0001** |
| *albidus – langeberg* | **<0.0001** | **<0.0001** | **<0.0001** | **<0.0001** | **0.0002** | 1.0000 | **0.0497** |
| *albidus – macowanianus* | **0.0109** | **<0.0001** | **<0.0001** | **<0.0001** | 0.5598 | **<0.0001** | **<0.0001** |
| *albidus – prismatosiphon* | **0.0005** | **<0.0001** | 0.0876 | **<0.0001** | 1.0000 | **<0.0001** | **0.0164** |
| *blandus – callistus* | 1.0000 | 1.0000 | **<0.0001** | **0.0303** | **<0.0001** | **0.0043** | 1.0000 |
| *blandus – high-altitude* | **0.0008** | 1.0000 | 1.0000 | 1.0000 | **<0.0001** | **<0.0001** | **0.0221** |
| *blandus – langeberg* | **<0.0001** | **0.0169** | 0.1135 | 0.1070 | **<0.0001** | **0.0033** | 1.0000 |
| *blandus – macowanianus* | 1.0000 | **0.0216** | **<0.0001** | **<0.0001** | 1.0000 | **<0.0001** | 0.4069 |
| *blandus – prismatosiphon* | **<0.0001** | 0.2758 | **<0.0001** | 0.0642 | **<0.0001** | 1.0000 | **0.0104** |
| *callistus – high-altitude* | **<0.0001** | 1.0000 | **<0.0001** | 0.9271 | **<0.0001** | **<0.0001** | **0.0017** |
| *callistus – langeberg* | **<0.0001** | 0.2683 | **0.0007** | 1.0000 | **<0.0001** | 1.0000 | 1.0000 |
| *callistus – macowanianus* | 0.0997 | 0.5099 | 0.0686 | 1.0000 | 1.0000 | **<0.0001** | **0.0175** |
| *callistus – prismatosiphon* | **<0.0001** | 1.0000 | 1.0000 | **<0.0001** | 1.0000 | **0.0286** | **0.0038** |
| *high-altitude – langeberg* | 0.3400 | **0.0103** | **0.0038** | 1.0000 | 1.0000 | **<0.0001** | 0.5813 |
| *high-altitude – macowanianus* | **0.0002** | **0.0110** | **<0.0001** | **0.0131** | **<0.0001** | **0.0015** | 0.3736 |
| *high-altitude – prismatosiphon* | 1.0000 | 0.2004 | **<0.0001** | **0.0014** | **<0.0001** | **<0.0001** | **<0.0001** |
| *langeberg – macowanianus* | **<0.0001** | 1.0000 | 0.0815 | 1.0000 | **<0.0001** | **0.0006** | 1.0000 |
| *langeberg – prismatosiphon* | 1.0000 | 1.0000 | 0.2197 | **<0.0001** | **<0.0001** | **0.0179** | **0.0001** |
| *macowanianus – prismatosiphon* | **<0.0001** | 1.0000 | 1.0000 | **<0.0001** | **0.0013** | **0.0179** | **<0.0001** |

**Table S5.** The mean and standard error of soil measurements taken at Jonaskop (JK), Kleinmond coast (KC), and Limietberg (LB).

| **Site** | **Potassium (mg/kg)** | **Sodium (mg/kg)** | **Phosphorus (mg/kg)** | **pH** | **Carbon (%)** | **$NH_4$ Nitrogen (%)** |
|---|---|---|---|---|---|---|
| JK | 28.00 ± 1.34 | 9.00 ± 0.63 | 13.60 ± 0.68 | 4.64 ± 0.02 | 0.71 ± 0.04 | 0.03 ± 0.01 |
| KC | 86.40 ± 2.93 | 283.60 ± 9.68 | 107.60 ± 5.07 | 5.52 ± 0.04 | 7.19 ± 0.21 | 0.41 ± 0.01 |
| LB | 95.20 ± 2.35 | 38.00 ± 3.26 | 18.60 ± 1.17 | 4.50 ± 0.00 | 0.62 ± 0.01 | 0.17 ± 0.01 |

**Table S6.** Pairwise comparisons between the soil measurements at Jonaskop (JK), Kleinmond coast (KC), and Limietberg (LB).

| **Comparisons** | **Potassium (mg/kg)** | **Sodium (mg/kg)** | **Phosphorus (mg/kg)** | **pH** | **Carbon (%)** | **$NH_4$ Nitrogen (%)** |
|---|---|---|---|---|---|---|
| JK - KC | **<0.0001** | **0.0006** | **0.0006** | **<0.0001** | **0.0591** | **<0.0001** |
| JK - LB | **<0.0001** | 0.1141 | 0.1320 | **0.0071** | 0.3409 | **<0.0001** |
| KC - LB | 0.058 | 0.1141 | 0.1048 | **<0.0001** | **0.0016** | **<0.0001** |

**Table S7**. Pairwise comparisons for the survival and height (mm) of Jonaskop (JK), Kleinmond Coast (KC), and Limietberg (LB) seedlings on each soil type. Soil type includes the native soil of all respective plant localities and a control soil. The control soil was used to test whether any ecotypes were suffering from transplant shock, therefore, only the survival is presented. Significant differences are highlighted in bold.

| Soil type | Ecotype comparison | Survival 2024 | Height 2024 (mm) |
|---|---|---|---|
| JK | JK - KC | 1.00 | **0.0441** |
| JK | JK - LB | 1.00 | 1.00 |
| JK | KC - LB | 1.00 | 0.31 |
| KC | JK - KC | 1.00 | 1.00 |
| KC | JK - LB | 1.00 | 1.00 |
| KC | KC - LB | 1.00 | 0.91 |
| LB | JK - KC | 1.00 | 1.00 |
| LB | JK - LB | 1.00 | 0.77 |
| LB | KC - LB | 1.00 | **0.0009** |
| Control | JK - KC | 1.00 | - |
| Control | JK - LB | 1.00 | - |
| Control | KC - LB | 1.00 | - |

**Table S8.** The survival rate (%) of all ecotypes at all four translocation sites in 2023 and 2024. All survival rates presented are the percentage of seedlings that survived from the beginning of the experiment.

| Site | Ecotype | Survival 2023 (%) | Survival 2024 (%) |
|---|---|---|---|
| JK | JK | 89 | 50 |
| JK | KC | 70 | 29 |
| JK | LB | 81 | 45 |
| KC | JK | 77 | 33 |
| KC | KC | 75 | 38 |
| KC | LB | 75 | 34 |
| LB | JK | 88 | 61 |
| LB | KC | 88 | 64 |
| LB | LB | 88 | 68 |

**Table S9**. Pairwise comparisons between Jonaskop (JK), Kleinmond Coast (KC), and Limietberg (LB) seedlings at all the reciprocal translocation sites. Generalized linear mixed models were used to test for differences in survival and height at the end of 2023 and 2024.

| **Site** | **Ecotype comparison** | **Survival 2023** | **Survival 2024** | **Height 2023 (mm)** | **Height 2024 (mm)** |
|---|---|---|---|---|---|
| JK | JK – KC | **0.0007** | **0.0028** | **0.0006** | 1.0000 |
| JK | JK – LB | 1.0000 | 1.0000 | 1.0000 | 1.0000 |
| JK | KC – LB | 0.5551 | 0.0760 | **0.0221** | 1.0000 |
| KC | JK – KC | 1.0000 | 1.0000 | **< 0.0001** | 1.0000 |
| KC | JK – LB | 1.0000 | 1.0000 | 1.0000 | 1.0000 |
| KC | KC – LB | 1.0000 | 1.0000 | **< 0.0001** | 0.0929 |
| LB | JK – KC | 1.0000 | 1.0000 | 0.1731 | 1.0000 |
| LB | JK – LB | 1.0000 | 1.0000 | 1.0000 | 0.3629 |
| LB | KC – LB | 1.0000 | 1.0000 | **0.0170** | **0.0033** |

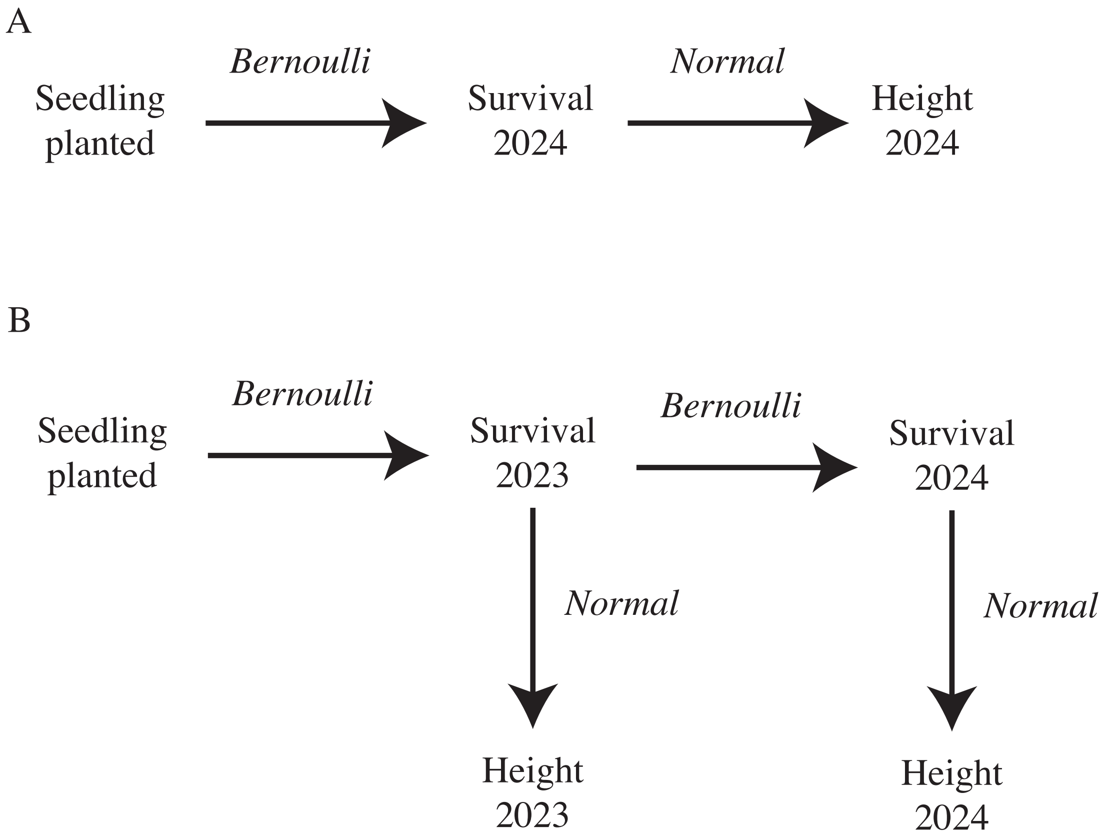


**Figure S1**. Aster model dependency structure used in the (A) common garden and (B) full 2023 and 2024 reciprocal translocation.
